# Chromosome-level genome assembly of the functionally extinct northern white rhinoceros (*Ceratotherium simum cottoni*)

**DOI:** 10.1101/2021.12.11.472206

**Authors:** Gaojianyong Wang, Björn Brändl, Christian Rohrandt, Karl Hong, Andy Pang, Joyce Lee, Harris A. Lewin, Giovanna Migliorelli, Mario Stanke, Remy Schwab, Sarah Ford, Iris Pollmann, Bernhard M. Schuldt, Marlys Houck, Oliver A. Ryder, Alexander Meissner, Jeanne F. Loring, Franz-Josef Müller, Marisa L. Korody

## Abstract

The northern white rhinoceros (NWR; *Ceratotherium simum cottoni*) is functionally extinct, with only two females remaining alive. Efforts to rescue the NWR have inspired the exploration of unconventional conservation methods, including the generation of artificial gametes from induced pluripotent stem cells and somatic cell nuclear transfer. To enable the technologies required for these approaches, we used complementary sequencing and mapping methods to generate a NWR chromosome-level reference genome that meets or exceeds the metrics proposed by the Vertebrate Genome Project. It represents 40 autosomes, an X and a partially-resolved Y chromosome, and the mitochondrial genome. We compared the NWR reference genome to the southern white rhinoceros (SWR) population that has been physically separated from the NWR for tens of thousands of years. Using short-read data from the SWR and optical mapping, we found that the two populations are very similar on both the chromosome level and mitochondrial genome level. The results of this study are encouraging for the efforts underway to rescue the NWR.

## Introduction

The northern white rhinoceros (NWR; *Ceratotherium simum cottoni*), with only two non-reproductive females alive, is functionally extinct due to human activities including poaching, civil war, and habitat loss and fragmentation [1–3]. The development of assisted reproductive technologies such as fertilization of harvested eggs with intracytoplasmic sperm injection, somatic cell nuclear transfer (SCNT), and generation of artificial gametes from induced pluripotent stem cells (iPSCs) may provide alternatives to save the NWR from extinction [4]. As part of an international plan to reestablish the NWR [4], iPSCs have been generated from the NWR and southern white rhinoceros (SWR) [5, 6] and embryonic stem cells derived from the SWR [7]. Based on genomic data from the NWRs, the iPSC lines are believed to encompass sufficient genomic diversity to reestablish a viable animal population [8]. The iPSCs would be differentiated into artificial gametes, which has been done in mouse [9–11], to generate embryos for implantation to surrogate mothers. A major challenge in developing these methods in NWR is the lack of a well-annotated reference genome. Such genomes have been foundational in human and mouse research for the understanding of stem cell biology and the development of directed differentiation methods.

Here, we report the genome assembly of a male NWR using long-read, short-read, and optical mapping technologies. The assembly is a 2.5 GB highly contiguous genome consisting of 40 partially haplo-phased chromosome-level autosomes with three chromosomal scaffolds (Chromosome 1, 11, and 23) potentially reaching telomere-to-telomere contiguity, a potential telomere-to-telomere X chromosome as well as a partially-resolved Y chromosome. The overall quality of the genome reaches or exceeds the metrics proposed by the Vertebrate Genome Project (VGP) [12]. Additionally, we report the long-read assembly of the NWR mitochondrial genome and provide evidence of consistent mitogenomic differences between the NWR and SWR populations based on short read data from 25 NWR and 27 SWR individuals.

The results obtained in this report not only provide an essential resource for unconventional conservation methods but also suggest that NWR and SWR populations are genetically distinct yet genomically compatible for genetic rescue via SCNT and potentially even hybridization.

### Assembly of the NWR reference genome

We obtained fibroblasts and established iPSC lines from a male NWR individual (“Angalifu”, laboratory number KB9947) [6]. The karyotyped NWR cells show 40 pairs of autosomes and one pair of allosomes (**Figure 1A**) consistent with published data [13]. We prepared, extracted and deeply sequenced high-molecular weight DNA from these cells using four technologies: Oxford Nanopore Technologies (ONT) long-reads with an average of 75X coverage, 10X Genomics linked reads (10XG) with an average of 80X coverage, Hi-C chromatin conformation capture with an average of 40X coverage, and Bionano optical mapping with an average of 400X coverage (**Materials and Methods)**.

**Figure 1.**
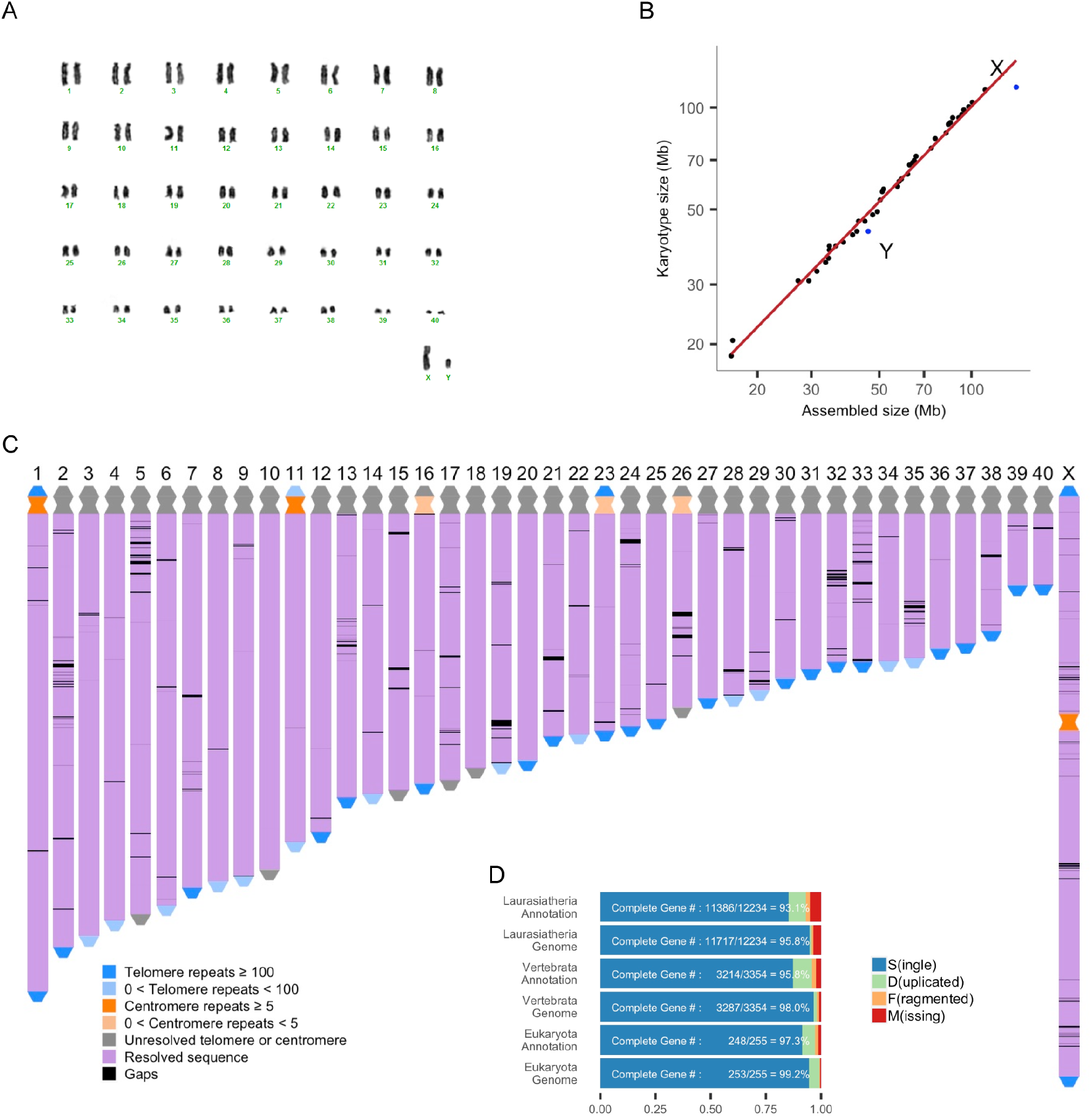
NWR genome assignment into chromosome-equivalent scaffolds. (A) giemsa-stained karyotype of “Angalifu” NWR 9947-c501 iPSCs, indicating a 2n=82 karyotype. (B) The assembled chromosomal level scaffold size in Mb compared to the estimated chromosomal size derived from the karyotype. (C) Visualization of the assembled NWR reference genome indicating resolved regions and gaps still remaining. (D) Benchmark Universal Single Copy Ortholog (BUSCO) scores for the genome assembly and the genome annotation using three datasets: Eukaryota (255 genes), Vertebrata (3354 genes), and Laurasiatheria (12234 genes).

#### Autosome assembly

We followed an automatic and manual assembly strategy similar to the workflow recently proposed by VGP [12] using ONT long reads rather than PacBio continuous long reads and customized the pipelines for allosome and mitochondrial genome assembly (**Supplementary Figure 1**, **Materials and Methods**). Contiguous sequences (contigs) were first assembled using the Shasta assembler [14] and ONT long reads. Scaffolds were polished using Racon [15] and Pilon [16] after three sequential rounds of scaffolding with 10XG linked reads, Bionano optical maps and Hi-C sequencing data obtained from the same individual (**Supplementary Figure 1A, Materials and Methods**). The first draft NWR genome assembly (fd) was generated after manual curation of the polished scaffolds (**Supplementary Figure 3A, B**) using Juicebox [17].

#### Allosome Assembly

We inspected the coverage profile of ONT reads aligned to fd (**Supplementary Figure 3C**). One assembled chromosome-level scaffold had a mean coverage of 38X, which is approximately half the 75X coverage of the other scaffolds and putatively corresponded to the X chromosome since the sequenced NWR individual “Angalifu” is a male. The remaining 40 assembled chromosome-level scaffolds putatively corresponded to the 40 autosomes of NWR. We found that some regions of the X chromosome in fd contained more coverage than expected, indicated by a localized increase in the coverage histogram (**Supplementary Figure 3C**). Due to the similarities between the X and Y in the pseudoautosomal region (PAR), and the repetitive sequences found at the boundaries of the PAR, a hybrid allosome scaffold containing X and Y sequences was assembled in fd.

Consequently, we re-assembled sequences corresponding to the Y chromosome with a customized pipeline (**Supplementary Figure 1B, Supplementary Figure 2**, **Materials and Methods**) by selecting out similar sequences of chromosome X and Y. Briefly, the re-assembled putative chromosome Y scaffolds were aligned back to the hybrid allosome scaffold in fd. The putative genomic sequences of chromosome Y that appeared to be mis-assembled into the hybrid allosome scaffold were removed based on the coverage profile. This resulted in the generation of a trimmed X chromosome (130.6 Mbps in total, updated coverage profile in **Supplementary Figure 3D**) and a partially-resolved Y chromosome (349 scaffolds, 41.2 Mbps in total). The reassembled chromosome-level X scaffold and Y scaffolds together with the previous 40 putative autosome scaffolds were called the second draft (sd) NWR genome assembly.

#### Mitochondrial Genome Assembly

We used a long-read baiting strategy to exclude nuclear mitochondrial DNA segments and to assemble the mitochondrial genome. The mitochondrial genome of the blue whale (*Balaenoptera musculus*) was used to select out all the mitochondrial reads from the ONT data to generate a preliminary assembly of 16,715 bps (**Supplementary Figure 1C, Materials and Methods**). We then curated the mitochondrial genome (mt), which was later added to sd to become the third draft (td) NWR genome assembly.

#### Curation

We curated td to close 307 small gaps (**Supplementary Figure 3E, Materials and Methods**) using ONT reads. We denoted the resulting genome assembly as CerSimCot1.0. It contains a complete mitochondrial genome as well as the chromosomal-level scaffolds (**Figure 1C**). We then named the autosomal scaffolds in CerSimCot1.0 as “assembled chromosomes” CHR 1 to 40 based on their sizes. The X chromosome scaffold in CerSimCot1.0 is named as CHR X and the remaining Y chromosome scaffolds are named CHR Y scaffold 1 to 349. We identified a 228bp NWR-specific centromeric repeat sequence from short reads [18] (**Materials and Methods**). Using this 228bp sequence as well as the telomeric repeat sequence (TTAGGG), we found that CerSimCot1.0 genome assembly has potentially achieved telomere to telomere assembly quality with CHR 1, 11, 23, and X, and centromere to telomere quality with CHR 16 (**Figure 1C**).

### Quality of NWR reference genome

The CerSimCot1.0 genome assembly meets or exceeds the metrics proposed by the Vertebrate Genome Project [12] (**Supplementary Table 1, Supplementary Figure 3H,**). The assembly resulted in a total of 2.50 Gbps with a contig NG50 of 3.6 Mbps and a chromosome-level scaffold NG50, which is within 8% of the estimated genome size of 2.69 Gbps (**Materials and Methods**). In each step of the genome assembly pipeline, there was an improvement in at least one of the VGP genome quality metrics (**Supplementary Figure 3F**). Overall, the accuracy of CerSimCot1.0 genome assembly has reliable blocks of 9.4 Mbps, phased blocks of 2.0 Mbps, base pair QV of 41.3 and k-mer completeness of 91.8% **(Supplementary Table 1**). Due to the complexity and larger amount of repetitive regions within the allosomes, particularly the Y chromosome, the assembly quality of CHR X and Y (QV range: 23 - 39: **Supplementary Figure 3G**) are lower than that of the CHR 1 - 40 (QV range: 34 - 44: **Supplementary Figure 3G**). In total, 427 gaps (approximately 160 gaps/Gbps) remain in the final genome assembly (**Supplementary Table 1**). Genome completeness was assessed using Benchmarking Universal Single-Copy Orthologs (BUSCO) [19] for three data sets: Eukaryote - 99.2% complete; Vertebrata - 98.0% complete; and Laurasiatheria - 95.8% complete (**Figure 1D, Supplementary Table 1**).

### Annotation of NWR reference genome

To annotate the genome, we first masked CerSimCot1.0 genome assembly (**Materials and Methods**), identifying approximately 33.22% of the genome as repetitive or low complexity, including 3.15% in short interspersed nuclear elements (SINE), 20.94% in long interspersed nuclear elements (LINE), 5.83% in long terminal repeats (LTR), 3.17% in DNA repeat elements (TcMar-Tigger and hAT-Charlie transposons). Genome and functional annotation were then performed using BRAKER1 [20] with a combination of RNAseq data from multiple tissues and cell types of NWR and SWR (**Supplementary Table 2**), followed by BRAKER2 [21] with five protein datasets (**Supplementary Table 3**), and TSEBRA [22] to combine the results from BRAKER1 and BRAKER2 (**Materials and Methods**). The BUSCO scores for the resulting genome annotation based on three datasets are: Eukaryota - 97.3% complete; Vertebrata - 95.9% complete; Laurasiatheria - 93.1% complete (**Figure 1D**).

### Chromosome conservation

The chromosomal-level scaffolds CHR 1 to 40, CHR X, and all the scaffolds of CHR Y agree well with the estimated chromosome sizes obtained from the karyotypes (**Figure 1B**). Cross-species chromosome painting experiments [23, 24] and preliminary fluorescent in situ hybridization (FISH) mapping studies [25] have reported a strong conservation of chromosome segments across the odd-toed ungulates (Order: *perissodactyla*) despite a range of karyotypes (2n = 32 – 2n = 84) [26]. To confirm this conservation at the genic level, we compared the domestic horse genome (*Equus caballus*, EquCab3.0, GenBank assembly accession: GCA_002863925.1 [27]) with the CerSimCot1.0 genome assembly (**Supplementary Figure 4A**). We successfully aligned all annotated genes in EquCab3.0 to CerSimCot1.0 (**Materials and Methods**), and observed a strong conservation of the gene order between horse and NWR (**Figure 2, Supplementary Figure 4**). Notably, we found that the genes located on opposite arms of horse Chromosomes 1, 2, 3, 6, 7, 8, 9, 10 and 15 mapped to separate chromosomes in NWR, which agrees with chromosome painting [24] and preliminary FISH mapping results [25].

**Figure 2.**
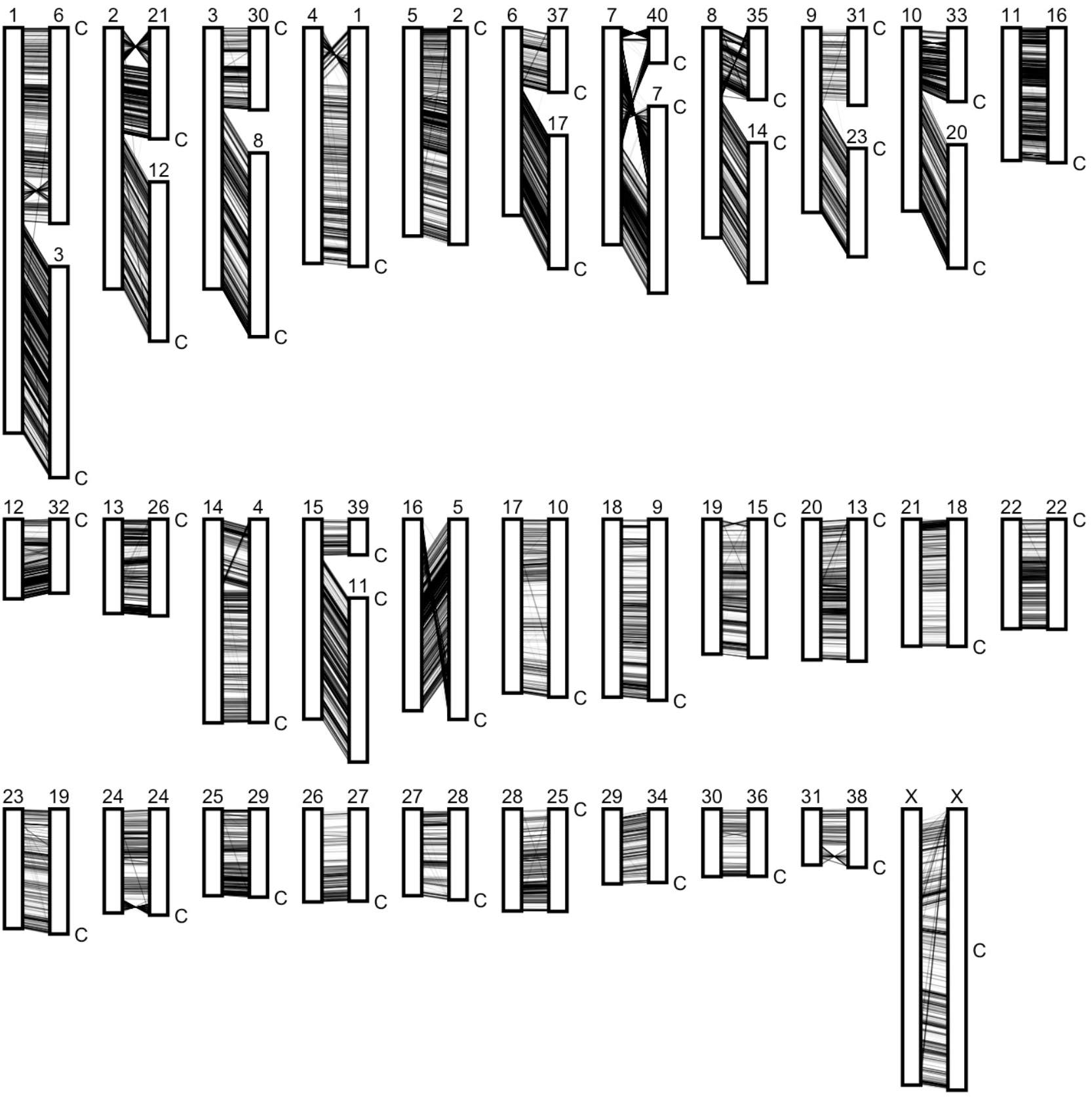
Gene order visualization between horse and NWR genomes represented at the chromosome level as karyotype ideograms. For each mapping relationship, left is the horse genome and the right is NWR genome. Chromosome arms and/or whole chromosomes are relatively conserved between the two species despite a divergence time of 52 – 58 million years [28].

### White rhino genomic comparison

To explore the genomic similarities and differences between NWR and SWR, we compared CerSimCot1.0 with the previously published SWR genome (CerSimSim1.0, GenBank assembly accession: GCA_000283155.1, **Materials and Methods**) and found several potential translocations between NWR and SWR (**Figure 3A**). Since assembly errors can be misinterpreted as translocations, we asked whether these apparent translocations were true genomic differences or instead assembly errors; we employed Bionano data from three SWR individuals (one male and two females) to re-scaffold the published SWR assembly with each of the individual-specific optical maps (**Materials and Methods**). Comparisons among these three re-scaffolded SWR genomes with the NWR CerSimCot1.0 genome assembly (**Materials and Methods**) demonstrated that the SWR and NWR appear to be strikingly similar at the chromosome level (**Figure 3B-D**), with no consistent large structural variations.

**Figure 3.**
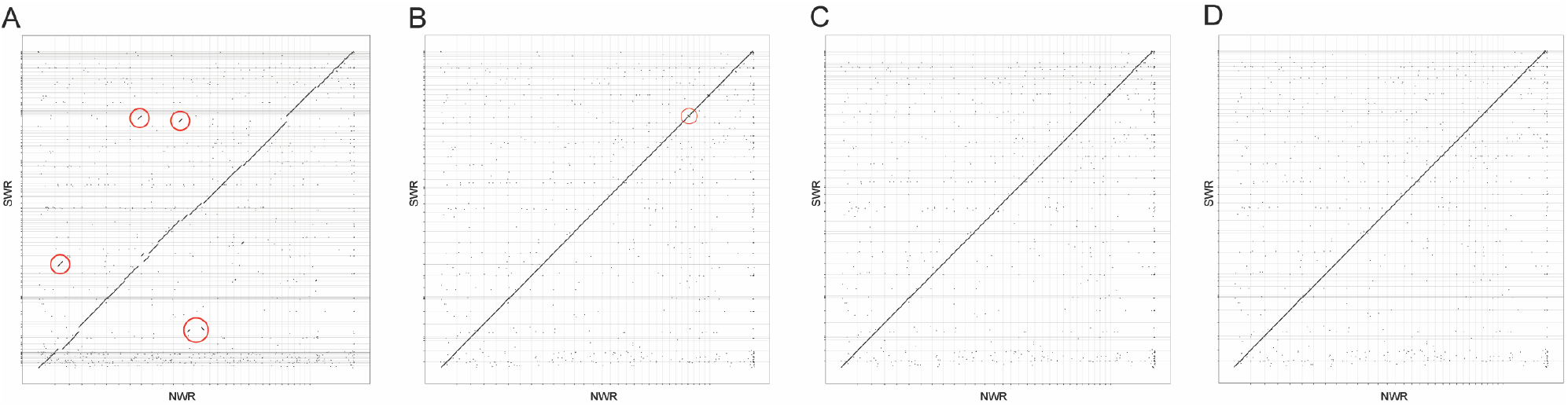
Genome comparison between NWR reference genome and (A) SWR genome CerSimSim1.0 (GCA_000283155.1). (B)-(D) Three SWR genomes scaffolded by optical mapping with the SWR genome CerSimSim1.0 (GCA_000283155.1) using SWR male KB10208 “Chuck” with a small translocation identified by the circle, SWR female KB 21328 “Amani”, and SWR female KB 21409 “Wallis”.

### Mitochondrial comparison of NWR and SWR

We explored the mitochondrial differences between NWR and SWR, with a focus on understanding the implications of their potential compatibility for SCNT with a SWR oocyte and NWR nucleus. We collected short read data of 25 NWRs and 27 SWRs from two different datasets (**Supplementary Table 4)** [8, 29], and identified consistent differences in SNPs between the SWR group and the NWR group. We identified 105 SNPs that differed between the SWR group and the NWR group (**Figure 4, Materials and Methods**), which is a Hamming distance of only 0.6%. At the protein level, five of the thirteen mitochondrial protein-coding genes contain consistent amino acid differences between SWR and NWR (**Supplementary Table 5**).

**Figure 4.**
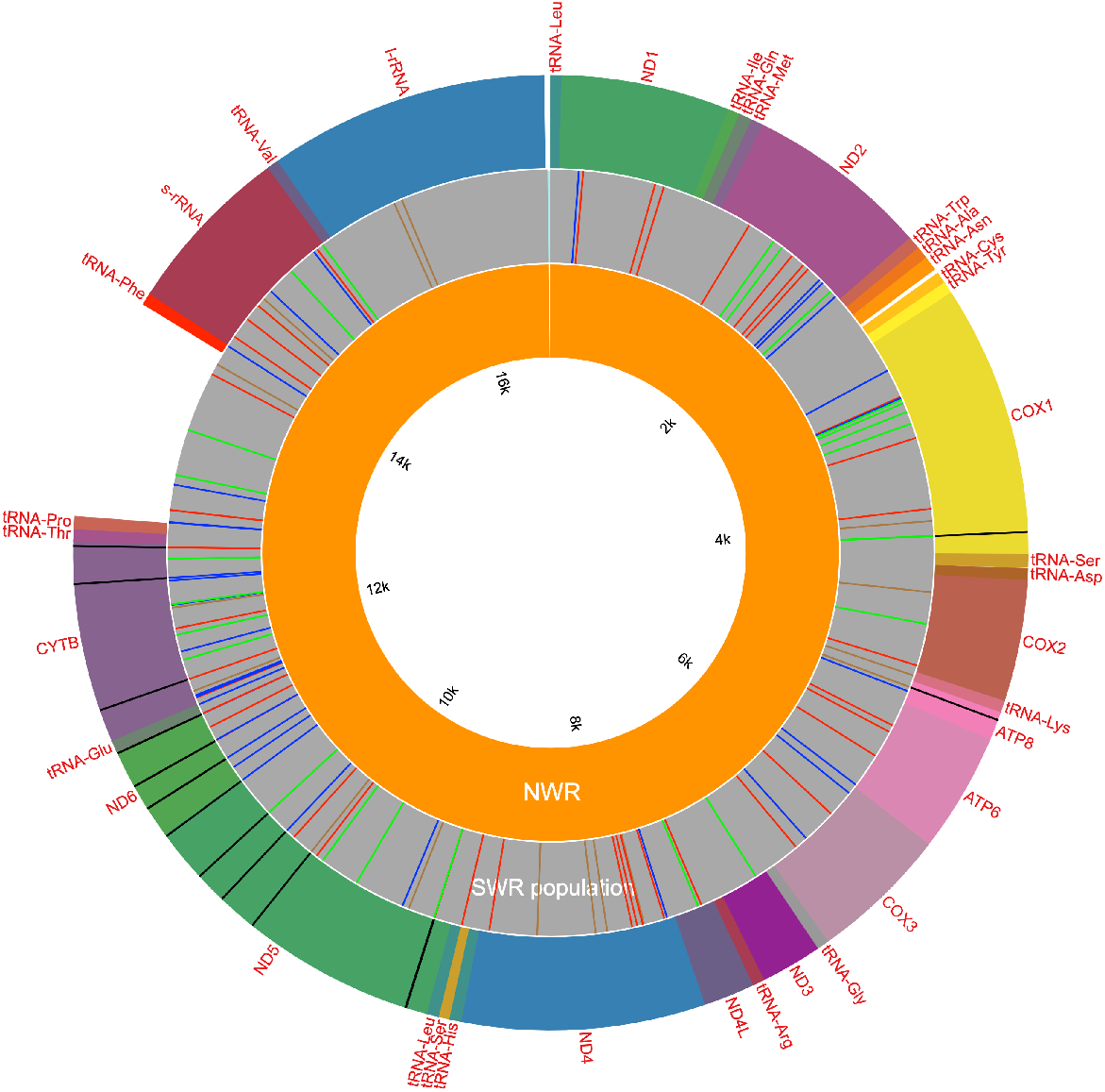
Mitochondrial genome comparison between NWR and SWR. Inner track (orange) is the NWR mitochondrial genome obtained from KB9947 “Angalifu”, the middle inner (gray) track is the population-level SWR mitochondrial genome with SNPs marked, the outer track is the localization of the 13 protein coding genes in the mitochondria with animo acid subistitutions marked in black.

## Discussion

Reference genome assemblies have become fundamental to scientific progress in biology. The NWR reference genome will enable the types of molecular genetic research in the white rhinoceros that have become standard for human and mouse stem cell scientists: identification of gene expression profiles of undifferentiated and differentiated pluripotent stem cells (PSCs), DNA methylation profiling, generation of reporter lines to develop directed PSC differentiation methods, gene targeting, and sequencing to detect genomic instability in PSCs. These methods will be essential for the generation of new NWR individuals using assisted reproduction technologies and artificial gametes.

NWR and SWR have been separated for tens of thousands of years [8, 30]. While the NWR population has declined to only two non-reproductive females, the SWR has been far more successful, becoming the most prevalent rhinoceros population after only approximately 100 individuals survived at the turn of the 20th century [31]. Genetic diversity was not a limiting factor in the SWR’s resurgence; interestingly, the genetic diversity of the SWR population as measured by heterozygosity surpasses that of humans [8]. For the NWR, there may be sufficient genetic diversity among the cryopreserved cells for re-establishing a sustainable population [8].

An international effort to genetically rescue the NWR [4] proposed a systematic list of approaches, all of which would require using SWR females as surrogate mothers. These approaches include SCNT using SWR oocytes as recipients of NWR nuclei, fertilization of oocytes recovered from the two females with NWR sperm, or generation of artificial gametes from the collection of iPSC lines from 9 NWR individuals [6, 9–11, 32].

The chromosomal number of SWRs and NWRs is identical, our analysis indicates that there are no large structural differences between their genomes, and we found no evidence to predict nuclear-mitochondrial incompatibilities. These findings reawaken the idea that fertile SWR/NWR hybrids might be possible. In 1977 a hybrid offspring (“Nasi” KB5767 SB #476) was born in captivity to a NWR mother (“Nasima” KB8174 SB # 352) and a SWR father (“Arthur” SB # 355) [33]. Nasi failed to reproduce during her 30-year lifespan, but this was not unusual; captive breeding of NWR and SWR was severely hampered by diet and management plans during that time. Based on a model using genetic distance as a proxy [34], our comparative analysis of the mitochondrial genomes of SWR and NWR predicts that hybrids such as Nasi could be fertile.

While generation of fertile SWR/NWR hybrids remains challenging and is an unexplored possibility, there are immediate practical applications of the NWR genome, such as assessing the feasibility of SCNT. In addition, reference genomes will aid in the assembly of genomes of other endangered rhinoceros and could be used in the future to measure relatedness in order to implement population-scale strategies such as managed breeding.

## Data availability

All raw data can be accessed through request to.

## Acknowledgements

The work would not be possible without the prescience and vision of Dr. Kurt Benirschke, who founded the Frozen Zoo in 1975. This work has been funded in part by the Germany Federal Ministry for Education and Research to AM, FJM, GW, and CR (IntraEpiGliom FKZ 13GW0347). AM, GW and RS are supported by the Max Planck Society.We acknowledge the support of David L. Barker for advice about available mapping technologies, the San Diego Zoo Wildlife Alliance and donors, especially Anne and Christopher Lewis, the Kleberg Foundation, the Alice B. Tyler Perpetual Trust, and the Weinberg Trust. We thank the IT department in MPIMG, particularly for the support from Thomas Kreitler.

## **Author contribution** (CRediT author statement)

GW: Conceptualization, Methodology, Software, Validation, Formal analysis, Investigation, Data Curation, Visualization, Writing - Original Draft, Writing – Review and Editing

BB, Methodology, Investigation, Writing – Original Draft

GM, MS: Methodology, Software, Data Curation

KH, AP, JL: Software, Resources, Investigation, Data Curation, Writing – Original Draft

CR, BS, RS: Methodology, Software, Data Curation, Writing – Original Draft

MH: Resources, Investigation

SF, IP: Investigation, Data Curation

HL, AM, OAR: Resources, Funding acquisition, Project administration

JFL: Methodology, Supervision, Writing – Review and Editing

FJM: Conceptualization, Investigation, Methodology, Validation, Resources, Visualization, Data Curation, Project Administration, Supervision, Funding acquisition, Writing - Original Draft, Writing – Review and Editing

MLK: Conceptualization, Investigation, Methodology, Resources, Visualization, Data Curation, Project Administration, Supervision, Funding acquisition, Writing - Original Draft, Writing – Review and Editing

## Supplementary Figures and Tables

**Supplementary Figure 1.**
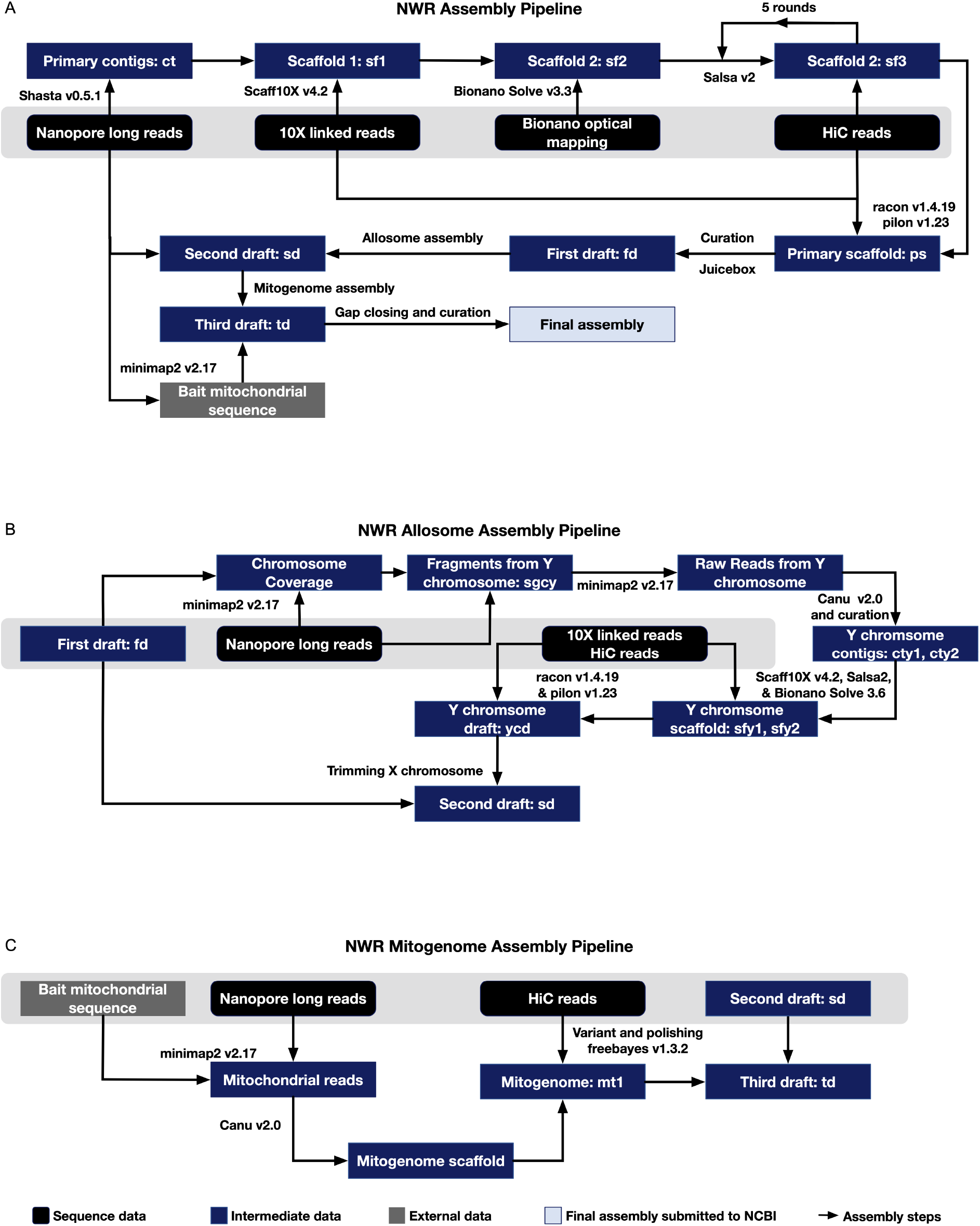
Flow charts of the assembly pipelines used for NWR genome. (A) Complete genome assembly pipeline used for the NWR reference genome: ONT long reads are assembled with Shasta into contigs. Scaffolds are generated with Scaff10x; are error corrected and scaffolded with Bionano Solve; and sacffolded with 5 rounds of Salsa with HiC data. Scaffolded results are polished and curated to create the first draft assembly (fd): fd is then used for the Allosome pipeline to create the second draft (sd): sd is then used for the mitochondria assembly pipeline to generate the third draft (td). (B) Allosome assembly pipeline: Y chromosome reads are selected from the ONT reads by overlapping with DNA sequences of hybrid allosome in fd with coverage ranging from 56 and 81, the selected reads are assembled using Canu; scaffolded with Scaff10X, Salsa, and Bionano; scaffolds are polished to generate the sd (C) Mitochondrial genome assembly pipeline: bait mitochondrial sequence from the blue whale was used to pull all mitochondrial reads from the ONT data; mitochondrial reads were then assembled using Canu; mitogenome (mt) was then polished and added to sd to become the td.

**Supplementary Figure 2.**
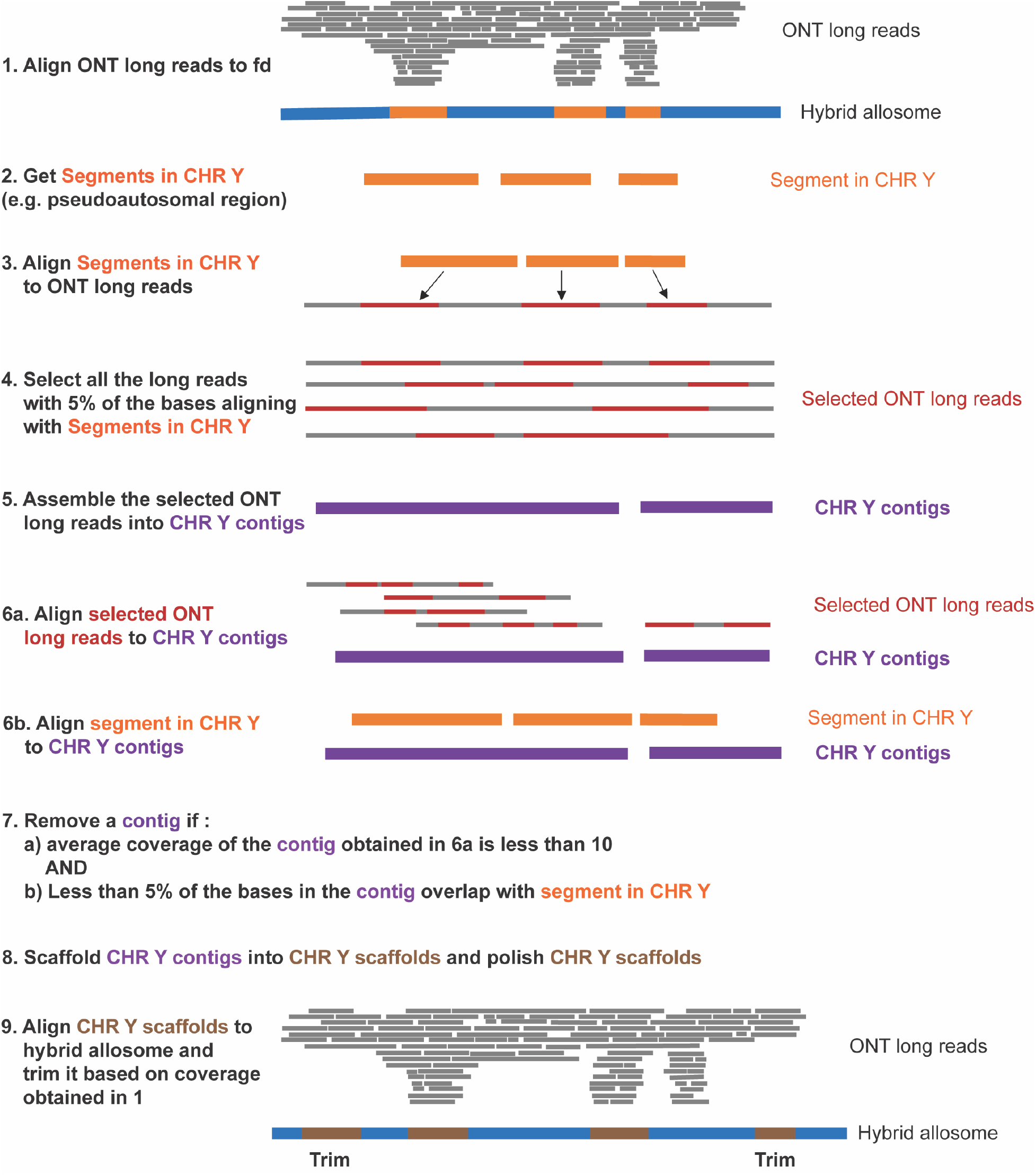
Detailed description of allosome assembly pipeline.

**Supplementary Figure 3.**
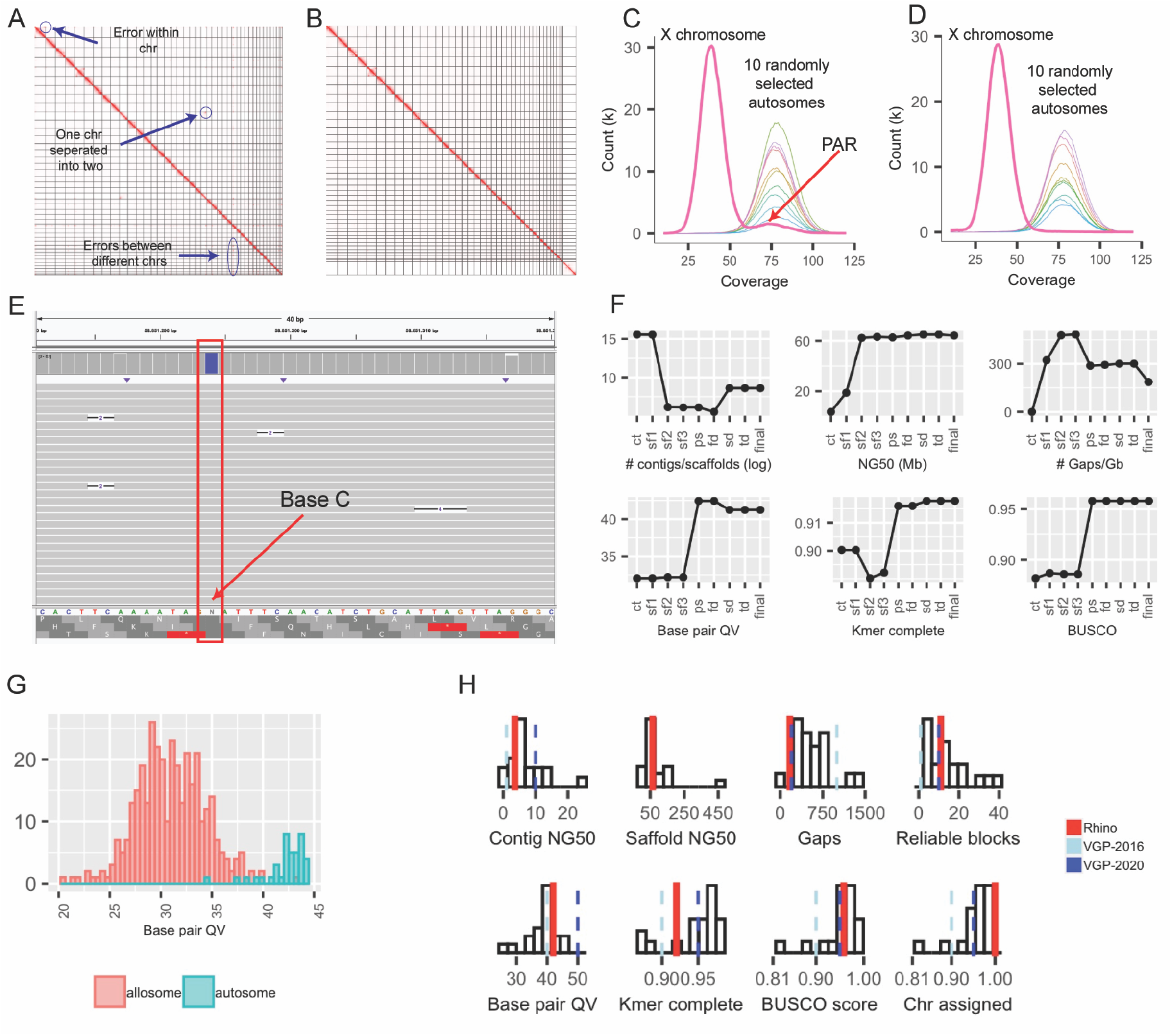
The improvement of assembly completeness and accuracy throughout the genome assembly pipeline and final assembly metrics. (A) The HiC contact map before any manual curation. (B) The HiC contact map after manual curation using Juicebox. (C) The coverage histogram (randomly sampled from 10 Mbps) of the X chromosome and 10 randomly selected autosomes in first draft (fd) (D) The coverage histogram (randomly sampled from 10 Mbps) of the X chromosome and 10 randomly selected autosomes in second draft (sd). (E) Using basecalling consensus from ONT reads to close gaps. (F) The improvements in genome quality at each metric throughout the assembly process. (G) The histograms of autosomes and allosomes base pair QV. (H) The quality of the NWR reference genome compared with the metrics proposed by the Vertebrate Genome Project, i.e. VGP-2016 and VGP-2020. The histograms represent the qualities of the 16 genomes generated by VGP [12].

**Supplementary Figure 4.**
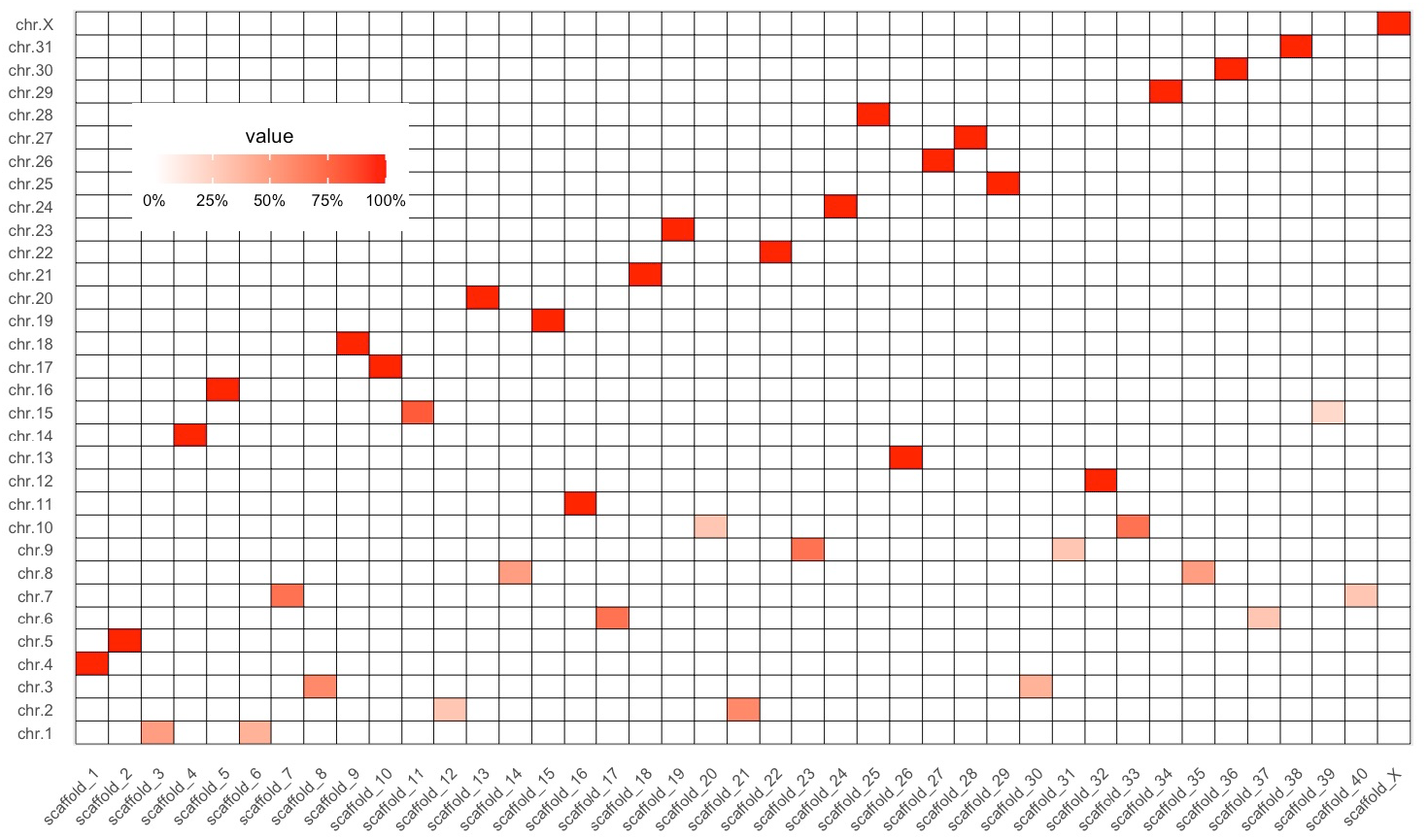
Conservation of chromosome segments and genes between the horse and NWR. The percent of horse genes in each chromosome aligned to NWR chromosomes is indicated.

**Supplementary Table 1:** Genome quality of NWR according to the proposed metrics of the Vertebrate Genome Project (part of the table comes from [12])

| Quality Category | Metric | Finished | VGP-2020 | VGP-2016 | VGP studies | NWR genome |
| --- | --- | --- | --- | --- | --- | --- |
| **Notation** | x.y.P.Q.C | c.c.Pc.Q60.C100 | 7.c.P6.Q50.C95 | 7.c.P6.Q50.C95 |  |  |
| **Continuity** | Contig NG50 (x) | =Chr. NG50 | >10 Mb | >10 Mb | 1–25 Mb | 3.6 Mbp |
|  | Scaffold NG50 (y) | =Chr. NG50 | =Chr. NG50 | =Chr. NG50 | 23–480 Mb | Chr. NG50 |
|  | Gaps per Gb | No gaps | <200 | <200 | 75–1,500 | 160 |
| **Structural accuracy** | Reliable blocks | =Chr. NG50 | >10 Mb | >10 Mb | 2.3–40.2 Mb | 0.2% |
|  | False duplications | 0% | <1% | <1% | 0.2–5.0% | 11 Mbp |
|  | Curation | Conflicts resolved | Manual | Manual | Manual | Automated + Manual |
| **Base accuracy** | Base pair QV (Q) | >60 | >50 | >50 | 39–43 | 42 |
|  | k-mer completeness | 100% complete | >95% | >95% | 87–98% | 92% |
| **Haplotype phasing** | Phase block NG50 (P) | =Chr. NG50 | >1 Mb | >1 Mb | 1.6 Mb<sup>a</sup> | 6.6 Mbp |
| **Functional completeness** | Genes | >98% complete | >95% complete | >95% complete | 82–98% | 95.9%<br>92.0% |
|  | Transcript mappability | >98% | >90% | >90% | 96% | NA |
| **Chromosome status** | Assigned (C) | >100% | >95% | >95% | 94.4–99.9% | 100% |
|  | Sex chromosomes | Right order, no gaps | Localized homo pairs | Localized homo pairs | At least one shared | Chr. X and partially Y resolved |
|  | Organelles (for example, MT) | One complete allele | One complete allele | One complete allele | One complete allele | One complete allele |

**Supplementary Table 2.** RNA-seq datasets used for annotation.

| **Supplementary Table 2. RNA-seq datasets used for annotation** |  |
| --- | --- |
| **Individual** | **Cell or Tissue Type** |
| NWR KB9947 | Fibroblasts |
| NWR KB9947 | Feeder-dependent iPSCs |
| NWR KB9947, KB8173, KB8175, KB6571, KB17626 | Embryoid bodies (14 days of differentiation) |
| NWR KB9947 | Embryoid bodies (28 days of differentiation) |
| NWR KB9947 | Embryoid bodies (38 days of differentiation) |
| NWR KB9947, KB8173, KB9939 | Embryoid bodies (Day 2 suspension, Day 4 suspension, Day 7 suspension, Day 2 plated, Day 4 plated, Day 7 plated) |
| NWR KB8173 and KB9939 | Incipient mesoderm-like cells – 60 hours of differentiation |
| NWR KB8173 | Primordial germ cell-like cells, Day 4 and Day 7 of differentiation |
| NWR KB6571 | Partially reprogrammed iPSCs |
| NWR KB8175 | Brain |
| NWR KB9947 | Adult Testes |
| SWR KB21638 | Neonate Testes |
| SWR NA | Mixed Female Reproductive Tissue |

**Supplementary Table 3:**
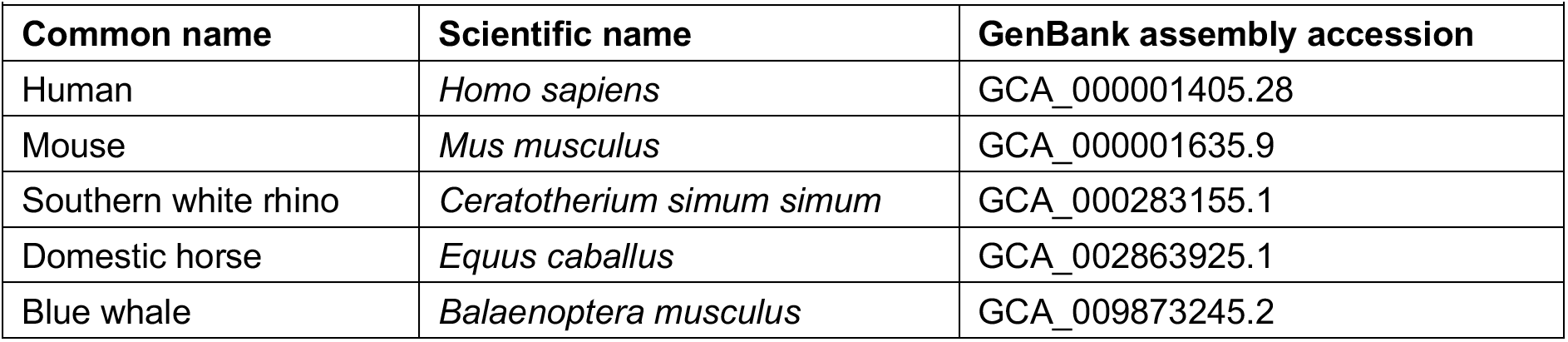
Protein datasets used to improve the annotation pipeline.

|  |  |  |
| --- | --- | --- |
| **Common name** | **Scientific name** | **GenBank assembly accession** |
| Human | *Homo sapiens* | GCA_000001405.28 |
| Mouse | *Mus musculus* | GCA_000001635.9 |
| Southern white rhino | *Ceratotherium simum simum* | GCA_000283155.1 |
| Domestic horse | *Equus caballus* | GCA_002863925.1 |
| Blue whale | *Balaenoptera musculus* | GCA_009873245.2 |

**Supplementary Table 4.** NWR and SWR short-read data.

| **Supplementary Table 4. NWR and SWR short-read data** |  |  |
| --- | --- | --- |
| **Individuals** | **Note** | **Accession** |
| KB3731, KB5763, KB5764, KB5766, KB7571, KB8174, KB8175, KB9939, KB9947, mwr_TG0450 ~ mwr_TG0456, mwr_TG0458, mwr_TG0460~ mwr_TG0463, mwr_TG0465, mwr_TGF14 ~ mwr_TGF16 | These samples are all NWRs | NCBI PRJNA74583, and European Nucleotide Archive (ENA) PRJEB45263 |
| KB5892, KB6974, KB7062, KB13306, KB21325, KB21326, KB21328, KB21329, KB21408, KB21409, KB21391, KB21423, KB21426, mwr_TG0449, mwr_TGF01 ~ mwr_TGF07, mwr_TGF09 ~ mwr_TGF13, mwr_TGF17 | These samples are all SWRs |  |

**Supplementary Table 5.** Differences of genes and transcripts in the mitochondrial genome between NWR and SWR populations. Protein-coding genes are red.

| Gene | NWR location | SWR location | #SNPs | # Amino acid substitutions in protein-coding genes |
| --- | --- | --- | --- | --- |
| tRNA-Leu | 4:62 | 4:62 | 0 | - |
| **ND1** | **65:1021** | **65:1021** | **4** | **0** |
| tRNA-Ile | 1021:1089 | 1021:1089 | 0 | - |
| tRNA-Gln | 1087:1159 | 1087:1159 | 0 | - |
| tRNA-Met | 1162:1230 | 1162:1230 | 0 | - |
| **ND2** | **1231:2274** | **1231:2274** | **10** | **0** |
| tRNA-Trp | 2273:2340 | 2273:2340 | 0 | - |
| tRNA-Ala | 2346:2414 | 2346:2414 | 0 | - |
| tRNA-Asn | 2416:2488 | 2416:2488 | 0 | - |
| tRNA-Cys | 2521:2586 | 2521:2586 | 0 | - |
| tRNA-Tyr | 2587:2653 | 2587:2653 | 0 | - |
| **COX1** | **2655:4199** | **2655:4199** | **12** | **1** |
| tRNA-Ser | 4197:4265 | 4197:4265 | 0 | - |
| tRNA-Asp | 4277:4343 | 4277:4343 | 0 | - |
| **COX2** | **4344:5027** | **4344:5027** | **3** | **0** |
| tRNA-Lys | 5031:5097 | 5031:5097 | 1 | - |
| **ATP8** | **5099:5305** | **5099:5305** | **2** | **1** |
| **ATP6** | **5260:5940** | **5260:5940** | **4** | **0** |
| **COX3** | **5940:6723** | **5940:6723** | **4** | **0** |
| tRNA-Gly | 6724:6792 | 6724:6792 | 0 | - |
| **ND3** | **6793:7138** | **6793:7138** | **1** | **0** |
| tRNA-Arg | 7140:7208 | 7140:7208 | 0 | - |
| **ND4L** | **7209:7505** | **7209:7505** | **2** | **0** |
| **ND4** | **7499:8876** | **7499:8876** | **10** | **0** |
| tRNA-His | 8877:8945 | 8877:8945 | 0 | - |
| tRNA-Ser | 8946:9004 | 8946:9004 | 1 | - |
| tRNA-Leu | 9006:9075 | 9006:9075 | 0 | - |
| **ND5** | **9076:10896** | **9076:10896** | **12** | **6** |
| **ND6** | **10880:11407** | **10880:11407** | **7** | **3** |
| tRNA-Glu | 11408:11476 | 11408:11476 | 1 | - |
| **CYTB** | **11481:12620** | **11481:12620** | **16** | **3** |
| tRNA-Thr | 12621:12689 | 12621:12689 | 0 | - |
| tRNA-Pro | 12691:12756 | 12691:12756 | 0 | - |
| tRNA-Phe | 14017:14085 | 14018:14086 | 0 | - |
| s-rRNA | 14086:15055 | 14087:15056 | 5 | - |
| tRNA-Val | 15056:15122 | 15057:15123 | 0 | - |
| l-rRNA | 15123:16700 | 15124:16701 | 2 | - |

