## Supplementary Materials for "Chromosome-level genome assembly of the functionally extinct northern white rhinoceros (*Ceratotherium simum cottoni*)"

### Materials and methods

#### *Sample Collection*

A wild-caught male northern white rhinoceros (NWR) was chosen as the reference individual: "Angalifu", lab ID KB9947 and studbook # 348. This individual was selected because it was previously identified [1] as a male with high runs of homozygosity relative to the NWR population to facilitate the assembly process. Fibroblast cell lines were obtained from the Frozen Zoo® part of the Wildlife Biodiversity Bank at the San Diego Zoo Wildlife Alliance and reprogrammed into induced pluripotent stem cells (iPSCs) as previously described [2]. These cells were then expanded, harvested, and cryopreserved as needed for each sequencing application.

#### *DNA Extraction, Genomic Library preparation, and sequencing*

*Oxford Nanopore Technologies (ONTs):* 30 – 50 x $10^6$ fibroblasts (passage 12) or iPSCs (passage 40) were harvested with 0.05% Trypsin - EDTA (Gibco) or Accutase (Corning) respectively, pelleted at 0.3g for 5 minutes, rinsed once with DPBS and resuspended in 100 μL of DPBS for phenol:chloroform extraction [3]. Briefly, cells were incubated for one hour at 37 °C in TBL buffer containing 10 mM Tris-CL, 25 mM EDTA, 0.5% SDS (W/V), and 20 μg/mL of RNase A. Cell suspension was then incubated for 3 hours at 50 °C with 100 μg/mL of Proteinase K, rotating end over end 10 times after each hour. Lysate was then poured into a tube containing phase-lock gel, and an equal volume of buffer-saturated phenol was added. Samples were rotated for 10 minutes at 40 rpm and centrifuged for 10 minutes at 4500 rpm. The aqueous upper layer was poured into a new phase-lock tube, and an equal volume of buffer-saturated phenol:chloroform was added. Samples were again rotated and centrifuged for 10 minutes. The aqueous upper layer was poured into a fresh tube, and a 1/10 volume of 7.5M ammonium acetate and 2.5 volumes of cold absolute EtOH were added. DNA was then spooled and rinsed in fresh 70% EtOH. Spooled DNA was washed twice with 80% EtOH and resuspended in 10 mM Tris-Cl pH 8 overnight. DNA concentration was then quantified in triplicate using a Qubit Fluorometer (Thermo Fisher Scientific).

30 μg of DNA was sheared using a Covaris ® g-Tube to 20 kbp by centrifuging for 60 seconds at 7,200 RPM. Alternatively, some DNA was sheared by passing through a 26g needle approximately 5 times. Sheared DNA was then size selected using a BluePippin (Sage Science) with either a 15 kbp or 20 kbp lid to remove small fragments.

Size selected DNA was then used for the Genomic DNA by Ligation (SQK-LSK109) library preparation (ONT, UK) according to the published protocol with minor adjustments: Up to 6 μg of DNA was used for each library preparation; DNA Repair and End-prep reaction incubation times were increased to 20 minutes at 20 °C and 10 minutes at 65 °C; End-prep reaction AMPure XP bead cleanup elution step was performed for 25 minutes at 60 °C with flicking every five minutes; Adaptor ligation was performed for 30 minutes at room temperature; AMPure XP beads were then incubated for 30 minutes at 48 °C with flicking and washed with "Long Fragment Buffer;" 80% EtOH was used throughout for all washing steps.

Sequencing was performed on a MinION instrument using R9.4.1 flow cells. Approximately 700ng of g-Tube sheared or 1.6 μg of needle sheared prepared library was loaded for each flow cell, and washes were performed after 20-24 hours using the Flow Cell Wash Kit (EXP-WSH003) and reloaded with a fresh library for a total run time of 72 hours.

*Bionano Genomics:* 1.5 x $10^6$ passage 8 fibroblasts were harvested using 0.05% Trypsin EDTA (Gibco), pelleted at 0.3g for 5 minutes, and resuspended in DMEM supplemented with 10% FBS and 10% DMSO. Cells were then slow-cooled in a Corning® CoolCell ™for at least 4 hours at -80 °C before continued short-term storage at -80 °C until processing.
Ultra-high molecular weight DNA was isolated using by using an agarose plug method (Bionano prep cell culture DNA isolation protocol #30026 revision F) or a silica disk protocol (Bionano Prep SP Frozen Cell Pellet DNA Isolation Protocol v2 #30398 revision A) to stabilize the DNA during isolation and purification. In short, DNA from 1.5 million cells was treated with RNase A and proteinase K in the presence of detergents and either embedded into an agarose plug or bound to a 4mm silica disk. The bound or embedded DNA was washed, eluted, and allowed to homogenize at room temperature overnight before fluorescent labeling. The following day, 750 ng was labeled at the recognition site CTTAAG using the enzyme DLE-1. The DNA was counter-stained and imaged on a Bionano Genomics Saphyr Gen II system until at least 400X raw coverage was collected for the genome assembly. At least 100X raw coverage was collected for structural variant detection.

*10X linked reads*: 1.5 x $10^6$ passage 6 fibroblasts were harvested using 0.05% Trypsin EDTA (Gibco), rinsed once with DPBS, and flash-frozen in liquid nitrogen. Frozen cells were then provided to the DNA Technologies and Expression Analysis Core at the UC Davis Genome Center (supported by NIH Shared Instrumentation Grant 1S10OD010786-01) for high molecular weight DNA extraction library preparation and sequencing.

*HiC sequencing:* 1.5 x $10^6$ passage 6 fibroblasts were harvested using 0.05% Trypsin EDTA (Gibco), rinsed once with DPBS, and flash-frozen in liquid nitrogen. Frozen cells were then provided to Dovetail™ Genomics (Scotts Valley, CA) for extraction, library preparation, and sequencing.

***RNA Extraction, Library Preparation, and Sequencing***

*RNA Extraction:* Cells were removed from the plate using either 0.05% Trypsin - EDTA (fibroblasts) or Accutase (iPSCs) as previously described, washed once in DPBS and resuspended in 600 μL of lysis buffer from *mir*Vana™ miRNA Isolation kit (Invitrogen) and stored at -80 °C until extraction. Tissue was collected and stored at -80 °C until use when it was finely minced on dry ice using a scalpel in the presence of lysis buffer. RNA was then extracted from the cells or tissue following the manufacturer's instructions. RNA was quantified using the Qubit fluorometer, and 100ng of total RNA was then used for the library preparation.

*Library Preparation:* Total RNA was Poly(A) selected using the NEBNext® Poly(A) mRNA Magnetic Isolation Module as part of the library generation protocol for the NEBNext® Ultra™ II Directional RNA Library Prep Kit for Illumina® (New England Biolabs, Ipswich, MA) following the manufacturer's instructions. RNA fragmentation was chemically performed at 94 °C for 15 minutes for a target insert size of 200 bp, and the PCR enrichment step was performed for 12 cycles. Library concentrations were determined using the Qubit, and insert sizes were obtained using the DNA High Sensitivity Bioanalyzer (Agilent Technologies) reaction and using these values diluted for sequencing. 76bp paired-end sequencing was performed on an Illumina NextSeq 500 by the Sanford Burnham Prebys Genomics Core (La Jolla, CA).

***NWR genome assembly***

*Contig Generation:* Guppy4 (ONT, UK) was used to base-call the ONT raw reads, and Shasta 0.5.1 [4] was used to generate the contigs from the ONT reads using parameters `--memoryMode filesystem --memoryBacking 2M –threads 128` for optimal performance. The contigs generated by Shasta are referred to as primary contigs (ct).

*10X scaffolding:* Scaff10X v4.2 [5] was used to compute an adjacency matrix after the 10X Genomic linked reads had been aligned to ct. One round of scaffolding was performed with the parameters `-nodes 63 -read-s1 12 -read-s2 8 -link-s1 10 -link-s2 10`. The generated scaffolds were referred to as scaffold1 (sf1).

*Bionano scaffolding:* Bionano Solve 3.3 (Bionano Genomics, Inc.) was utilized to scaffold sf1 with the haplotype-unaware optical genome mapping (OGM) molecules obtained from "Angalifu". The scaffolding process was performed with parameters: pre-assembly ON, extend and split OFF, cut complex multi-path regions ON. The scaffolds generated with the Bionano optical maps were referred to as scaffold2 (sf2).

*HiC scaffolding:* The mapping pipeline from Arima Genomics [6] was used for the alignment of Hi-C reads to the scaffold sf2. In the mapping pipeline, both ends of a read pair were individually aligned to sf2 using BWA-MEM [7] with default parameters. Only the 5'-side of the chimeric reads mate-pair were retained, and the 3'-side were removed from the two bam files generated by BWA-MEM in the previous step. Then a sorted and filtered bam file of paired single-end HiC reads was generated from combining the two bam files and filtering out the reads with mapping quality (MAPQ) less than 10. Picard Tools [8] were used to remove PCR duplicates existing in the pair-end bam file, the last step of the Arima Genomics mapping pipeline. Salsa2 [9] was used to estimate the orientation and location of contigs based on the normalized frequency of HiC interactions between them. Five rounds of scaffolding were performed with the capability to identify misassemblies in sf2 using parameters `-m yes -i 5 -p yes`. The restriction enzyme for Arima HiC data is set by parameter `-e GATC`. After 5 rounds of scaffolding, the generated scaffolds were referred to as scaffold3 (sf3).

*Polishing*: One round of Racon polishing [10] and one round of Pilon polishing [11] were performed using HiC reads and 10X linked reads. Racon was performed with default parameters using bam files generated from the HiC reads and 10X linked reads aligned to sf3. Pilon was performed with parameter `java -Xmx512G -jar pilon-1.23.jar` using bam files generated from the HiC reads and 10X linked reads aligned to the polished scaffolds from Racon using BWR-MEM [7]. The ONT reads were aligned to the polished scaffold from pilon, and all scaffolds with mean coverage lower than 40 or higher than 200 were removed. The scaffolds after polishing and scaffolds removal were called polished scaffolds (ps).

*Juicebox curation*: We then applied the Juicer [12] and 3D-DNA [13] pipeline to ps. We then manually curated ps using Juicebox [14], correcting minor errors in ps, and combined ps into 41 scaffolds, referred to as the first draft of the rhino assembly (fd).

*Allosome assembly:* We aligned all the nanopore raw reads to fd using minimap2 [15] and obtained the coverage profile of each scaffold. The coverage histogram of one scaffold, corresponding to a hybrid allosome, contains two peaks centered at 38 and 75 in contrast to the coverage histograms of the remaining 40 scaffolds, corresponding to the autosomes, containing only one peak centered at 75 (**Error! Reference source not found.**A). Since only one copy of chromosome X and one copy of chromosome Y exist, those regions in the hybrid allsome with the same coverage compared with the autosomes are regions present in both X and Y chromosomes.

We identified all the bases in the hybrid allosome with coverage values between 56 and 81. Continuous DNA sequences bounded by any selected bases with a distance less than 50 bps were selected out. All the selected DNA sequences longer than 100 bps were called segments in CHR Y (sgcy). Sgcy is aligned to each nanopore read. In order to find the raw reads probably coming from chromosome Y, we selected out all the raw reads with more than 5% bases overlapping with sgcy. Canu was used for the assembly of the selected reads, referred to as CHR Y contigs (cty1).

We reasoned that the raw reads selection process described above could not remove all the raw reads from chromosome X. These raw reads from chromosome X would probably end up being unassembled or assembled with low coverage in Canu [16]. Even if some raw reads from chromosome X are able to be assembled into few contigs with high coverage, these few contigs would not be similar to sgcy at all. We then 1) align the selected reads sgcy to cty1 and 2) align sgcy to cty1. We removed a contig in cty1 if this contig meets the following two criteria: 1) average coverage of the contig is less than 10 and 2) less than 5% of the bases in the contig overlap with sgcy. We generated contigs of a total size of 46 Mbps, referred to as cty2.

Cty2 was scaffolded by 10X linked reads [5], and HiC sequencing [6], similar to the scaffolding of the whole nuclear genome. The generated scaffold is called sfy1. Then sfy1 was scaffolded using Bionano Solve 3.6 (Bionano Genomics, Inc.) by aligning sfy1 to the Bionano OGM molecules. The algorithm iteratively extended the NGS contigs using Bionano's long molecules and merged any overlapping contigs resulting from the extended length. The generated scaffold from Bionano is called sfy2. Then sfy2 was polished once using Racon [10] and Pilon [11] to generate CHR Y scaffolds (ys).

We aligned ys back to the hybrid sex chromosome. The regions where the hybrid sex chromosome overlaps with ys should be either DNA sequences in both chromosomes X and Y or DNA sequences in chromosome Y but are misassembled into chromosome X. The DNA sequences that exist in chromosome Y but are misassembled into chromosome X should have a coverage half of the coverage in autosomes. Therefore, we trimmed the hybrid sex chromosome where it overlaps with ys and with low coverage (less than 38) to generate an updated version of chromosome X. The updated X and Y chromosomes and the autosomes in the first draft were merged to generate the second draft of the rhino assembly (sd).

*Mitogenome assembly:* All the ONT raw reads were aligned to the mitogenome [17] of *Balaenoptera musculus* (blue whale) using minimap2 [15]. Since a mitogenome is usually less than 19kb, those reads whose length was less than 19kb and whose MAPQ was larger than 30 were selected. These selected reads were assembled into a contig of 33kb using Canu [16]. The front of this contig was identical to its rear, suggesting it should represent a circular genome, which was then manually trimmed to remove the overlap. After manual trimming, a contig of 16,715 bp was obtained, representing the complete mitogenome of "Angalifu". The mitochondrial genome was then corrected by ONT reads through manual curation. The mitogenome (called mt) was merged to sd to generated the third draft of the rhino assembly (td).

*Gaps and manual curation:* We aligned all the ONT reads to td using minimap2 [15] and curated the alignment results in Integrative Genomics Viewer [18]. The ONT reads covering both ends of a gap were used to close the gap. The final assembly after the curation of td was called the final draft of the rhino assembly.

***Genome size estimation***

We used Meryl [19] to count 31-mers from the ONT raw reads and generate histograms from these reads. Genome size, repeat content, and heterozygosity were then estimated using GenomeScope [20].

***Metrics of genome assembly quality:***

Continuity: 1) The contig/scaffold NG50, defined as the size of a contig/scaffold where the sum of all the contigs/scaffolds with sizes longer than this scaffold is larger than half of the estimated genome size, is the current standard in the measure of genome assembly continuity. The northern white rhino assembly scaffolds have reached a Chr.NG50, which represents the highest standard in genome assembly. 2) Gaps, given by the number of gaps existing in the final assembly, quantify the number of unresolved regions in the genome.

Base accuracy: 1) Base pair QV was estimated by a reference-free method integrated into the Merqury pipeline [19]. Since k-mers uniquely identified in the assembly and not shown in the raw data are caused by errors, the assembly k-mers only existing in the genome assembly can be used for base-pair QV estimation. 2) K-mer completeness, obtained using the Merqury pipeline [19], is given by the k-mer found in an assembly divided by the k-mers found in the raw read data.

Structure accuracy: 1) False duplications: If a genome is completely sequenced with a coverage of *c*, then a unique k-mer from a homozygous (diploid) or heterozygous (haploid) region can be found *c* or *c/2* times. Any extra k-mers are caused by false duplication. We used Merqury [19] to count the number of distinct k-mers in the genome with additional copies and estimated the ratio of false duplications. 2) Reliable blocks were given by the concordance of at least two types of sequencing data out of four (ONT, 10X linked reads, Bionano, and HiC) support the assembled structure at each base [21].

Haplophasing: We aligned the pair-end HiC reads to the final draft of the northern white rhino genome using BWA-MEM [7] with default parameters. Freebayes [22] was used to call variants from the HiC-short-read bam file with coverage below 30 (i.e.: -g 30). We aligned the ONT raw reads to the final draft of the northern white rhino genome using minimap2 [15]. Whatshap [23] was used to phase the identified variants and to generate haplophased blocks using the generated ONT-long-read bam file with default parameters.

Functional completeness: BUSCO (Benchmarking Universal Single-Copy Orthologs) [24] V5.1.2 was used to evaluate the functional completeness of the generated assemblies. Assemblies were compared with lineage datasets eukaryota_odb10 containing 255 genes, vertebrata_odb10 containing 3554 genes, and laurasiatheria_odb10 containing 12234 genes.

*Telomere and centromere assesment:* The telomere sequence (TTAGGG) is conserved across various species [25]. To identify the centromere sequence of northern white rhino, we downloaded the Illumina short-read WGS sequencing data (SRR11428438) of a northern white rhino from NCBI. We randomly sampled 100k reads from this dataset and assembled the sampled reads into contigs using PRICE (Paired-Read Iterative Contig Extension) assembler [26]. The repeat sequences in the generated contigs were identified using TRF (Tandem Repeat Finder) [27]. This resulted in the identification of a 228bp repeated sequence:

TTTGGCTCAGTTCTGGGAATCTTTGAACGTCCTATACAAACTTGTTCCTAGATAG
TAACTTGCAGCTTGGAATGAGAAAGTGCTTGCTTACAGCCAAGTGCTGTATGTT
CTGAATGCCCGCATTGGGAATCTGTGTCAAAACTTTGTTAGGAATTGTGTGATCC
TTACTAGCTGGAAGTAAGGGAGATGGGGCATGAAGTTATTTCTAGGAGTTGGAG
ACATACCCAC

***Mapping of horse genes to NWR genome***

All horse genes were aligned to the NWR genome using BWA-MEM [7], and the coordinates of these alignment results were extracted. We used a scoring matrix (each row represents a horse chromosome and each column represents an NWR chromosome) to trace the gene mapping relationships between horse and NWR: 1) a horse gene from any horse chromosome had one point, and 2) this point was distributed equally to the NWR chromosome where this gene was aligned to. We then normalized the scoring matrix by row and obtained the percentage of genes on each horse chromosome aligned to the NWR chromosomes.

#### *Chromosome assignment*

The X and Y chromosomes have been defined in previous sections. Consecutive integer numbers were assigned to the remaining autosomes based on their sizes, i.e., the longest autosome is chromosome 1 while the shortest autosome is chromosome 40.

#### *Softmasking genome*

The final draft genome was soft masked using RepeatMasker [28] with the command `RepeatMasker -pa 15 -qq -species mammal -gff -xsmall -nolow`

#### *Genome and functional annotation*

The RNA libraries were aligned to the NWR assembly using STAR aligner [29]. We ran BRAKER1 [30] using the generated alignment files with pre-trained AGUSTUS parameters of human, i.e. `--species=human --softmasking –skipAllTraining`, and ran BRAKER2 [31] using protein datasets with pre-trained AGUSTUS parameters of human, i.e. `--prg gth --skipAllTraining --species=human –-softmasking`. The results from BRAKER1 and BRAKER2 were fixed, merged, and renamed using TSEBRA [32]. All the predicted genes or transcripts were blasted and functionally annotated against the Swiss-Prot dataset from the UniProt Knowledgebase [33].

#### *Comparison between NWR and SWR genomes*

The generated CerSimCot1.0 reference genome and the SWR reference genome (CerSimSim1.0, GenBank assembly accession: GCA_000283155.1) were compared using Minimap2 [15] with parameters `-x asm5`. The generated results were visualized using dotPlotly [34] with parameters `–m 2000 -q 500000 -l -p 12`. Several translocations and inversions were identified between CerSimCot1.0 and CerSimSim1.0. In order to determine whether these structural variants were due to assembly errors or represented true biological differences, we used Bionano optical mapping to scaffold DNA from three SWRs with CerSimSim1.0.

***Scaffolding of SWR genome***

Hybrid scaffolding of the SWR genome assembly (CerSimSim1.0, GenBank assembly accession: GCA_000283155.1) was performed using the "resolve all conflicts" parameter, which resolves conflicting alignments between CerSimSim1.0 scaffolds and optical genome maps by assessing molecule support at the conflicting region. If the optical genome map was supported by molecules that span the conflicting region, then the CerSimSim1.0 scaffold was cut and re-scaffolded. If the optical genome map does not have molecule support that spans the conflicting region, then the map was cut and re-scaffolded with the CerSimSim1.0 scaffold (Bionano Solve Theory of Operation: Hybrid Scaffold #30073 revision F).

***Comparison between NWR and the scaffolded SWR genomes***

The generated CerSimCot1.0 reference genome and the scaffolded SWR reference genomes were compared using Minimap2 [15] with parameters `-x asm5` and visualized using dotPlotly [34] with parameters `-m 2000 -q 500000 -l -p 12`.

***SNP calling of mitogenome***

The short-read data of an NWR or an SWR individual was aligned to mt using BWA-MEM [7], and the average coverage of mt (ave_cov_mt) was obtained. In order to remove the influence of nuclear mitochondrial pseudogenes, SNPs were called using Freebayes [22] with parameter `-C "$($ ave_cov_mt /3)"`, which means a SNP is only considered if it is supported by number of reads higher than 1/3 of the average coverage of mt.
